# Comprehensive identification of bacterial and archaeal habitat associations using IndicPiper

**DOI:** 10.64898/2026.11.19.767919

**Authors:** Clifton P. Bueno de Mesquita, Ben J. Woodcroft, Noah Fierer

## Abstract

Taxa that are prevalent and abundant in a particular habitat more so than other habitats can be considered indicators of that habitat. In addition to providing basic natural history information, indicator taxa can be used to verify metadata assignments in large public databases, identify potential contamination, and assess contributions of different source populations to a sink population. Despite these potential applications, a comprehensive database of microbial indicator taxa for major habitats of interest does not yet exist. Here we present IndicPiper, a database and methodology that uses the Sandpiper database combined with indicator taxa analysis to identify the habitat associations of bacterial and archaeal genera. IndicPiper includes a clearly defined methodology and set of functions such that any user with a list of habitats of interest could run the functions and generate their own database. IndicPiper uses the Genome Taxonomy Database (GTDB) for taxonomy, and the provided database or custom databases can be used to analyze any prokaryotic sample-by-taxa abundance table, including those from shotgun metagenomic and 16S rRNA marker gene sequencing. After presenting the methodology and results, we then present three applications of IndicPiper on actual modern and ancient DNA sequencing datasets. We used IndicPiper to assess the contribution of different source habitats to Lake Tjörnin sediments in Iceland over the past 2000 years, to paired indoor and outdoor dust samples from homes across the U.S, and to freshwater samples across the Willamette River watershed in Oregon, USA.

## Introduction

Indicator taxa are defined as the most characteristic taxa of a given group of samples, found mostly in a single group and present in the majority of samples belonging to that group [1, 2]. Quantitative indices of indicator taxa are maximized when a taxon is only found in a single group (e.g., habitat, site, treatment etc.) and when the taxon occurs in all samples of that group at high relative abundances [1]. For example, the commonly used IndVal metric takes into account the mean relative abundance of a taxon in one group compared to all groups (specificity), as well as the proportion of samples in which that taxon is observed in that one group (sensitivity) [1]. The identification of indicator taxa is already a widespread analysis in both macroecology and microbial ecology, with applications for monitoring, conservation, and management [3]. Habitat indicator taxa are taxa that are mostly found in a specific habitat and are found in many samples of that habitat type, while taxa that are generalists or are found more evenly in two or more habitats are simply considered “non-indicators”.

Here we developed a tractable and generalizable methodology (IndicPiper) that we then used to generate a comprehensive database of bacterial and archaeal habitat indicator genera. We envision IndicPiper serving three main purposes. First, such a database provides basic ecological information about those ‘indicator’ genera. Currently, it is relatively straightforward to obtain information about the isolation source of particular microbial genomes (e.g., available on the National Center for Biotechnology Information database or Genome Taxonomy Database), but this information cannot necessarily be used to identify if that same taxon could also be found in other habitat types. It is more informative to know how a taxon is distributed than where it was isolated from or the particular characteristics of the environment from which the genome was assembled. Given a particular genus of interest in a given study (i.e., a highly abundant and/or prevalent one, or a differentially abundant one), it would be useful to know if that genus is associated with a particular habitat to aid in interpretation of results.

Second, for certain sample types, such as air, ancient deposits, or other environments considered “sinks”, where taxa immigrate in from other environments, the database would be a useful and tractable way to identify the likely source habitats of the microbial taxa that end up in the sink environment. While this can be done qualitatively by searching the literature to identify putative habitat associations, there are pre-existing tools to do this in a more robust, quantitative, and consistent manner. For example, SourceTracker [4] and meta-SourceTracker [5] are methods that calculate the proportion of a given sample that is sourced from a given community.

However, these methods are highly dependent on the subset of samples that are selected as the potential source environments and use a Bayesian approach that is a ‘black box’ to most users. If the set of input samples is not representative of the environment type, then the results will be biased. IndicPiper uses a fundamentally different approach than SourceTracker in that it uses a pre-generated database made with all of the available metagenomic profiles on Sandpiper [6]. Alternatively, prior source-tracking research has used either a similar approach as IndicPiper (indicator taxa analysis on a set of samples from different habitats), but with a more limited number of input samples [7], or a combination of indicator taxa analysis and literature review [8]. IndicPiper does not use literature review but expands the indicator taxa analysis to be much more comprehensive by utilizing the Sandpiper database and by performing iterations over multiple random subsets of samples from each habitat.

Third, such a database of habitat indicator genera could also be useful for identifying contaminants, when taxa from different habitats are inadvertently mixed (e.g. skin bacteria being introduced into marine water samples), for verifying or adding metadata assignments in public databases, or more generally as an initial data quality check. Lists of potential contaminants exist [9, 10], but must be used carefully as they include taxa that could also be truly present in a sample and these lists of contaminants do not necessarily include taxa that could be introduced by inadvertent mixing of sample types during collection or processing. IndicPiper simply provides information on the habitat associations of taxa, which users can take into account when considering whether samples may or may not be contaminated. Furthermore, one of the first things that researchers do after generating data on microbial community composition is to ask if the taxa in their samples make sense, generally relying on knowledge of the literature. IndicPiper offers an additional tool and more formalized approach for performing such a check, as researchers could use IndicPiper to determine if the dominant taxa are expected in a given sample type.

We present the development and application of IndicPiper, a tool for generating lists of habitat indicator genera using the comprehensive Sandpiper database [6], which as of May 2026 (version 2.0.1) contains information on which bacterial and archaeal taxa are detected in over 900,000 metagenomes. We describe the methodology and results of the IndicPiper analyses as well as three use cases that demonstrate how IndicPiper can be applied to different scenarios in microbial ecology to aid in interpretation of microbial community sequencing data from shotgun metagenomes or 16S rRNA gene amplicons.

## Materials and Methods

### Inputs and Filtering

We used a combination of SingleM taxonomic profiles publicly available in the Sandpiper database [6] and indicator taxa analysis [1] to develop a list of genera associated with different habitats. Sandpiper (https://sandpiper.qut.edu.au/) v2.1.0 contains SingleM profiles based on GTDB [11] and GlobDB [12] genomes and taxonomy for 913,165 publicly available metagenomes. We first downloaded the publicly available Sandpiper sample metadata (“parsed_metadata” and “per_acc_summary” files), the additional sample metadata generated by Kingfisher [13], and the Sandpiper GTDB r232 coverage table from Zenodo (https://zenodo.org/records/20437114). The BioSample accession, study accession, and BioProject accession from Kingfisher were merged with the Sandpiper metadata table. The coverage table was converted to a site-by-taxon relative abundance table at the genus level using the SingleM “summarise” function.

We performed a variety of database filtering steps prior to performing the indicator analysis, which are outlined in the filterSandpiper.R script. Samples were kept if they were real bulk metagenomes with no cell sorting or single cell techniques used, had a minimum of 150 million archaeal and bacterial base pairs as estimated via SingleM prokaryotic fraction [14], had no warning flags, passed the low complexity filtering flag, had at least 40% relative abundance assigned to genus level in their SingleM profile, and had a clear and specific habitat designation in the “organism” column (derived from NCBI metadata) in the Sandpiper metadata (e.g., “human gut metagenome” was retained but “gut metagenome” was removed). We also removed duplicate sequencing runs that were from the same BioSample, such that each BioSample was only represented once. Lastly, we removed samples from habitats represented by less than 56 samples per habitat.

Next, we subset the genus relative abundance table to each individual habitat and calculated the Bray-Curtis dissimilarity of all pairwise sample comparisons within each habitat. To remove extreme outliers that could be from misclassified samples due to erroneous user-provided metadata, we filtered out samples with mean within-habitat Bray-Curtis dissimilarity values > 3 standard deviations above the mean. This resulted in the removal of 2209 samples (Table S1). The result of all filtering steps was a final metadata file (n = 426,084 samples) and genus relative abundance file (n = 37,188 genera), with a minimum of 52 samples per habitat. These files are publicly available on Zenodo (https://zenodo.org/records/23424117) and are the starting points for the IndicPiper pipeline (see below).

### IndicPiper Pipeline

There are three main functions to IndicPiper (Figure 1): prepIndicPiper, runIndicPiper, and checkIndicPiper. First, prepIndicPiper filters the provided starting metadata table and genus relative abundance table to a user-defined list of habitats of interest. Users can see a list of available habitats and their sample sizes with the utility function countHabitats. Users also decide in prepIndicPiper whether to lump or split certain habitats that are expected to share many bacterial and archaeal genera: mammalian gut samples (human, pig, bovine, sheep, canine, goat), rhizosphere and soil samples, ice and glacier samples, human oral and saliva samples, and different freshwater samples and labels (aquatic, freshwater, pond, lake water, riverine). In our example analysis (described below), we opted to combine these habitats since they share many genera, we were more interested in comparisons of broad habitat types, and none of those specific habitats were relevant to our research questions. In practice, users will have a choice whether to combine these habitats or keep them separate at the prepIndicPiper step based on their samples and study goals. In general, we recommend combining these habitats unless one of those habitats is specifically under investigation. The filtered metadata and genus relative abundance data are then saved and used as inputs to runIndicPiper.

**Figure 1.**
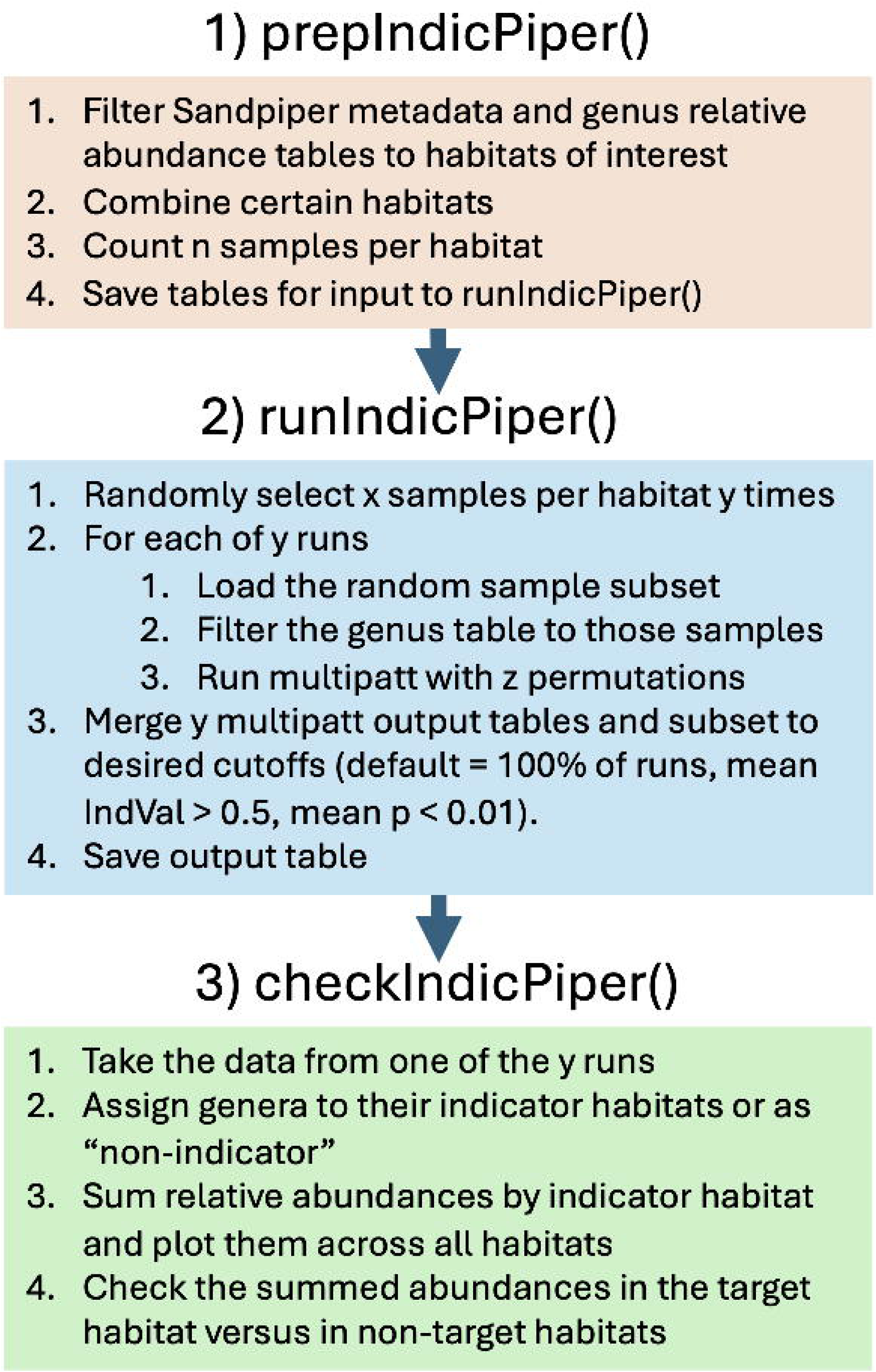
Overview of the three main functions of IndicPiper for generating a database of the habitat associations of bacterial and archaeal genera.

Second, runIndicPiper performs study-accession-stratified random subsampling of the habitats to the same sample size, indicator taxa analysis with the multipatt function in the IndicSpecies R package [2] over multiple iterations, and merges and filters the multipatt outputs from all of the iterations with user-defined thresholds. Multipatt is run using the “IndVal.g” metric [1, 2] and 100 permutations by default to calculate significance. The IndVal.g index is defined as the square root of specificity times fidelity, where specificity is the probability that a sampled site belongs to group j given that species i was observed there, adjusted so that groups are weighted equally regardless of their sample sizes, and fidelity is the proportion of sites in group j where species i occurs. The specificity component is based on the mean abundance of species i in group j relative to its mean abundances across all groups [1, 2].

The random subsampling approach is balanced partitioning with cyclic reuse. This approach balances the sample size per habitat and is also stratified by study accession to ensure no large study dominates any of the iterations. For habitats with more samples than the number of samples to draw per habitat times the number of iterations, each sample is used only once, and not all samples are used. For other habitats, samples must appear in multiple iterations, and the degree of repetition is greater in habitats with smaller sample sizes (Table S2). In those habitats, the iterations are not completely independent due to the reuse of samples. The iterative approach enables testing of the consistency of indicator taxa over multiple random subsets of each habitat, and the percent of iterations in which a genus is a significant indicator of the same habitat is one of the filtering criteria (“run_cut” argument). Users provide the number of samples per habitat to subsample per multipatt iteration as well as the number of multipatt iterations; we recommend setting the number of samples to slightly below the lowest number of samples per habitat, and 100 iterations of multipatt to balance speed with significance testing. Computational time scales linearly with the number of multipatt iterations. Users also define the number of permutations used by multipatt in each iteration to determine significance (100 by default). Users should ensure their p-value cutoffs are not less than the number of permutations used by multipatt (with 100 permutations, the minimum p-value cutoff should be 0.01, which is the default). For the thresholds to filter the combined multipatt outputs and identify significant indicator genera of each habitat, we set default parameters as follows: genus is present in at least one sample in all 100 multipatt iterations, is indicative of the same habitat in 100% of multipatt iterations, and has a mean p-value < 0.01 and mean IndVal.g of > 0.5 across the multipatt iterations.

The most important cutoff is the mean IndVal.g, which defines the strength of the association. We suggest a cutoff of 0.5 and set this as a default to balance having relatively strong indicators while capturing more taxa than using a more stringent cutoff. We tested cutoffs of 0.5, 0.6, and 0.7, and found that the number of indicators identified and their mean summed abundances in their indicator habitats decreased, with only minor improvements to mean summed relative abundances in non-indicator habitats (Table S3, Figure S1, Figure S2). Users can increase the IndVal cutoff if they want to identify fewer but stronger indicators, but we recommend using 0.5 as the minimum threshold to be confident in calling a genus an indicator.

Lastly, checkIndicPiper uses the output of one of the iterations to sum the relative abundances of indicator genera by habitat and then plot the relative abundances across all habitats, which enables users to see what percent abundance of the community is captured by the indicator genera and to ensure that relative abundances are higher in the indicated (“target”) habitats compared to all of the other (“non-target”) habitats (Table 1, Figure 2). The function checkIndicPiper also prints the mean and median summed relative abundances of the indictor genera in the target and non-target habitats. IndicPiper is publicly available on GitHub (https://github.com/cliffbueno/IndicPiper), and releases are synced with Zenodo for stable DOI tracking (IndicPiper v2.1.0 is available at https://zenodo.org/records/23481965).

**Table 1.**
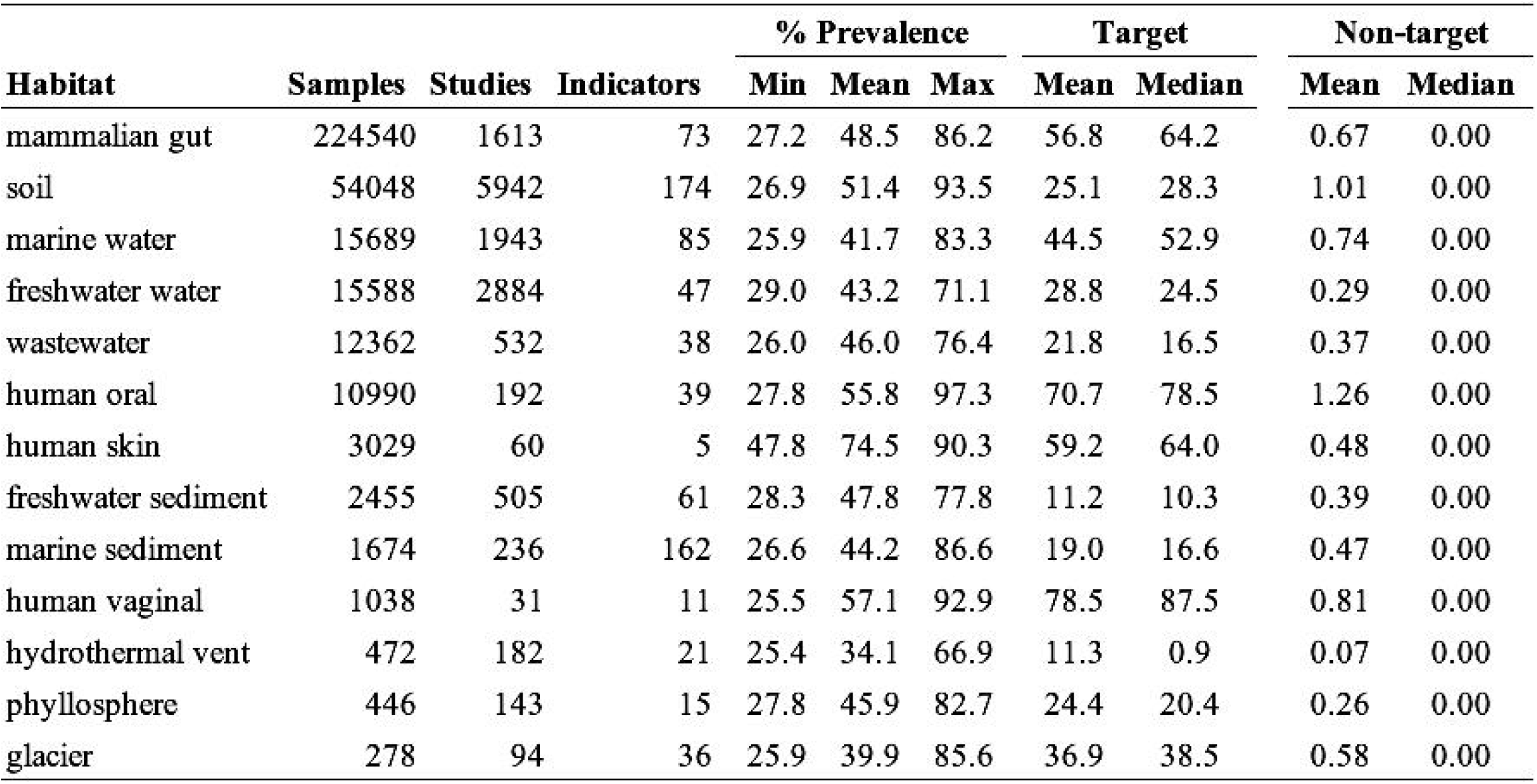
Number of samples, studies, and indicator genera for each of the 13 habitats for IndicPiper v2.1.0. The number of studies per habitat was calculated using the “study_accession” column in the Kingfisher metadata file for Sandpiper v2.0.1 (https://zenodo.org/records/20437114). The minimum, mean, and maximum percent prevalence of individual indicator genera across all samples of their target habitat (the habitat of which the genera are indicators) are shown. Also shown are the mean and median summed relative abundances of indicator genera in the target habitat and the non-target habitats (the other 12 habitats), for one of the random sample subsets (n = 225 per habitat).

**Figure 2.**
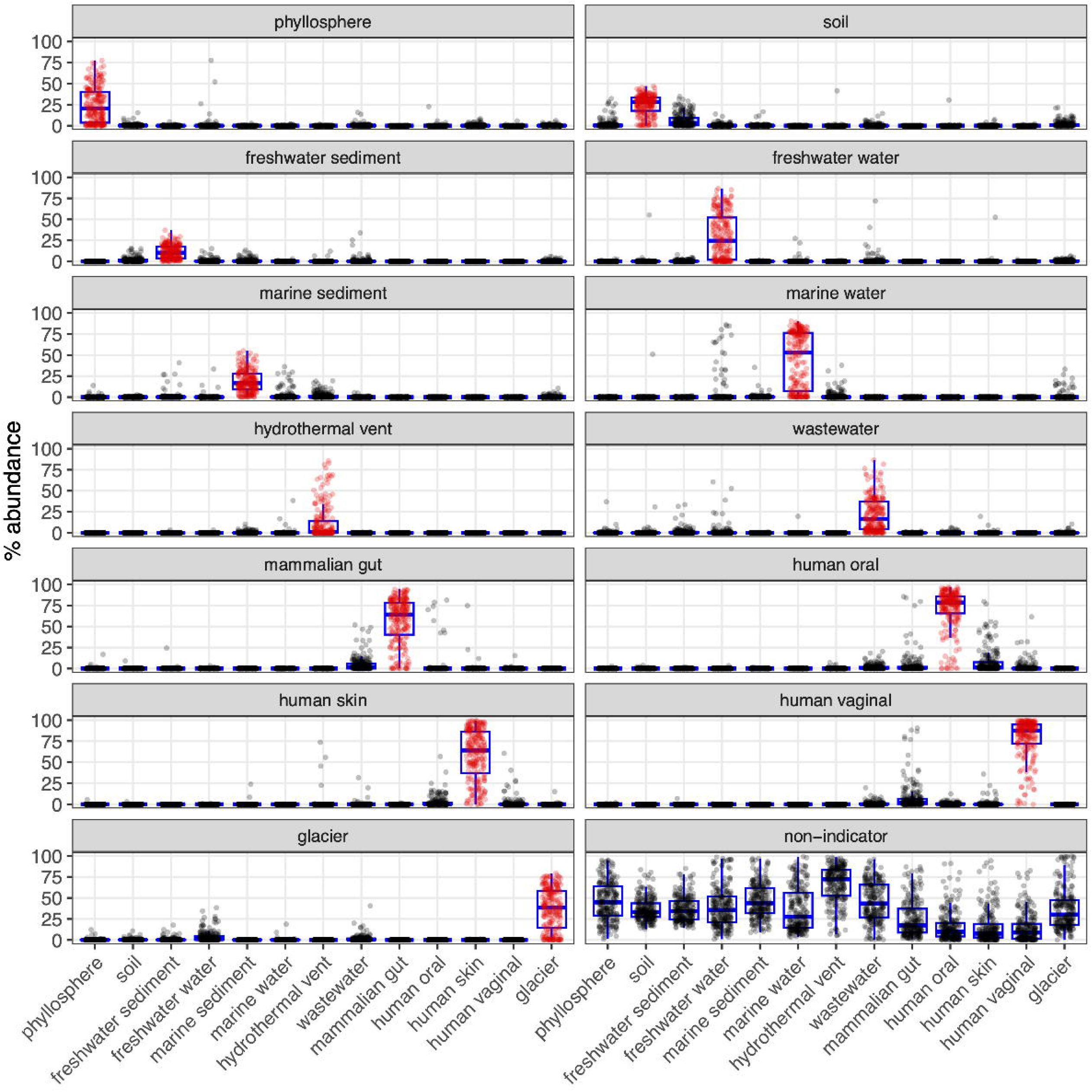
Summed relative abundances of indicator genera of each given habitat across all habitats for the publicly available IndicPiper v2.0.1 database we generated with prepIndicPiper() and runIndicPiper(). n = 225 metagenomes per habitat (one of the 100 random subsets of data). The figure was made with checkIndicPiper(). Target habitats (the habitat of which the genus is an indicator) are shown in red. While there is some spillover, habitat indicator genera typically sum to much higher values in target habitats than in non-target habitats (see Table 1 for exact means and medians). The “non-indicator” panel shows the summed relative abundances per sample of all genera that were not classified as indicators by IndicPiper.

### Provided IndicPiper Database

We used the starting files and the three IndicPiper functions to generate a publicly available IndicPiper database for 13 environmental and host-associated habitats with broad applicability to ecological studies: phyllosphere, soil, freshwater sediment, freshwater water, marine sediment, marine water, hydrothermal vent, wastewater, mammalian gut, human oral, human skin, human vaginal, and glacier. Sample numbers per habitat ranged from 278 (glacier) to 224,540 (mammalian gut), while the number of study accessions per habitat ranged from 31 to 5942 (Table 1). We ran prepIndicPiper providing a list of the 13 habitats and setting all of the combine sample arguments to true. The exact text strings used at this step are defined in Table S4. We then ran runIndicPiper with 225 samples per habitat, 100 multipatt iterations, and the default indicator cutoffs described above (100% run consistency, mean p-value < 0.01, mean IndVal > 0.5). Since mammalian gut and soil habitats had > 22,500 samples, there was no repeated sampling in any of the iterations for those two habitats, while for the other 11 habitats there was necessarily some repeated sampling (Table S2). The degree that samples had to appear in multiple runs is directly related to the sample size, such that glacier had the most samples repeated across iterations (Table S2). We then ran checkIndicPiper to plot the summed abundances of the indicator genera across each habitat for one of the iterations (Figure 2). The computational resources used to run each step of the pipeline are presented in Table S5.

The product of the analysis is a database for public use that contains the complete GTDB taxonomy, the associated habitat, the percent of runs that the genus was a significant indicator of the same habitat, and the minimum, maximum, mean, and standard errors of the IndVal and p-values from the runs. The table is publicly available on the GitHub repository (https://github.com/cliffbueno/IndicPiper, file = genus_habitat_indicators_v2.1.0.csv) and Zenodo (https://zenodo.org/records/23481965). A tabular version of the indicator species list, sorted by habitat and mean IndVal, is also available in the Supplementary Material (Table S6). Any researcher with a standard prokaryotic sample-by-taxonomy (GTDB taxonomy) abundance table can analyze the relative abundance of the habitat indicator genera, either individually or summed by habitat, by exact matching of genus names.

### Case Studies

To demonstrate the utility of the IndicPiper database, we conducted three case studies using publicly available data. In case study one, we analyzed ancient environmental DNA samples from Lake Tjörnin sediments in Iceland [15], calculating the aggregated relative abundance of indicators of different habitats to understand different microbial inputs to the lake over time. Shotgun metagenomes were processed as detailed in Kjær et al. (2025), with taxonomy assigned using a custom alignment-based approach optimized for ancient DNA [16]. Since GTDB r226 was used in their analysis, we used IndicPiper v1.0.0 for that analysis, which used Sandpiper v1.0.1, which contained GTDB r226 profiles from ∼700,000 metagenomes. The main difference between IndicPiper v1.0.0 and v2.0.0 is the Sandpiper version (and consequently GTDB version) that was used as input data.

For the other two case studies, we used IndicPiper v2.1.0 with GTDB r232 taxonomy. In case study two, we analyzed dust samples from inside and outside of homes from across the U.S. [17], investigating differences in indicator taxa present in the indoor versus outdoor dust. We reanalyzed the publicly available sample-by-taxa table that was generated with 16S rRNA marker gene sequencing data and the UPARSE pipeline [18], and reassigned taxonomy using the assignTaxonomy function in the DADA2 R package [19] with the sativa-curated GTDB r232 16S rRNA gene taxonomic database (https://figshare.scilifelab.se/articles/dataset/SBDI_Sativa_curated_16S_GTDB_database/14869077) for the indicator analysis, and with SILVA v138.2 [20] to remove sequences assigned as chloroplasts, mitochondria, or eukaryotes. Sequences lacking a domain-level assignment were removed, as were singletons and doubletons. Samples with < 1200 reads were excluded, and sequence counts in the remaining samples were converted to relative abundances. Differences in the summed relative abundances of indicator genera between indoor and outdoor dust samples were tested with paired t-tests, with each household containing one indoor sample and one outdoor sample. Only households with one of each sampling location (indoor, outdoor) were retained in the analysis (n = 1276 households).

In case study three, we analyzed 16S rRNA gene sequencing data from across the Willamette watershed in Oregon, USA [21]. We downloaded the raw sequencing data from 69 samples across the watershed, processed them with the DADA2 pipeline to quality filter sequences, infer amplicon sequence variants (ASVs), and assign taxonomy with the GTDB r232 database as above. Chloroplast, mitochondria, and eukaryote ASVs were removed based on classification with the SILVA v138.2 database. ASVs lacking a domain-level assignment were removed, as were singletons and doubletons. We rarefied the data to 5348 sequences per sample, which removed 6 of the 69 samples. We then filtered out replicates from the same exact geographic coordinates, such that each sample was from a unique location (n = 38). We categorized the samples into three categories - mountains, plains, urban - based on their geographic location. Differences in the summed relative abundances of indicator genera among the three categories were tested with ANOVA followed by Tukey’s Honest Significant Difference post hoc test.

## Results

We filtered the Sandpiper v2.0.1 database to 13 habitats of interest, with a minimum sample size of 278 and a maximum sample size of 224,540, and a minimum number of unique studies per habitat of 31 (Table 1). After running IndicPiper with the default parameters (described above), we identified a total of 767 indicator genera across those habitats (Table S6). Human skin had the fewest indicator genera with only five, while soil had the most indicator genera, with 174 (Table 1). These 767 indicator genera were distributed across 40 phyla, with different phyla represented in different habitats, and the number of genera per phylum ranging from 1 to 66 (Figure 3).

**Figure 3.**
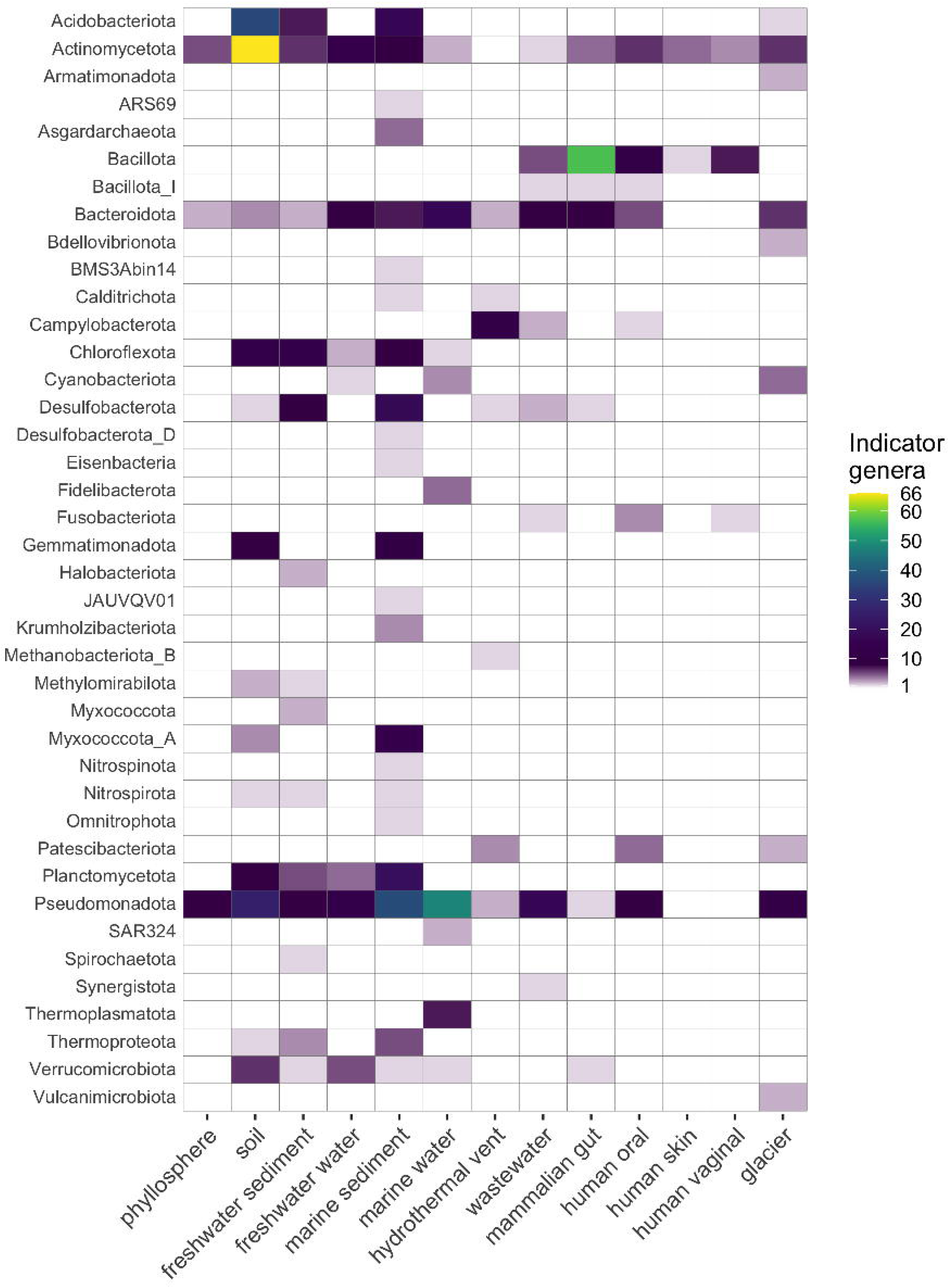
Number of indicator genera per phylum per habitat.

The top (highest indicator values) indicator genera of each habitat were generally in line with the literature on those habitats. The top five indicators per habitat were: freshwater sediment - *Bathycorpusculum*, *LD21, CAIXSE01*, *CTSoil-043*, *Methanoregula*; freshwater water - *Polynucleobacter*, *Planktophila*, *Limnohabitans_A*, *Limnohabitans*, *Sediminibacterium*; glacier - *Polaromonas*, *Lacisediminihabitans*, *Ferruginibacter*, *Rhodoglobus*, *Sphingomonas_O*; human oral - *Actinomyces*, *Rothia*, *Pauljensenia*, *Streptococcus*, *Granulicatella*; human skin - *Cutibacterium*, *Staphylococcus*, *Corynebacterium*, *Lawsonella*, *Micrococcus*; human vaginal - *Bifidobacterium*, *Lactobacillus*, *Fannyhessea*, *Berryella*, *Megasphaera*; hydrothermal vent - *Sulfurimonas, Sulfurovum*, *UBA1140*, *Sulfidibacterium*, *Thermococcus*; mammalian gut - *Blautia_A*, *Mediterraneibacter*, *Roseburia*, *Phocaeicola*, *Faecalibacterium*; marine sediment - *UBA1847*, *JACZVU01*, *SG8-38*, *Nitrosomaritimum*, *SZUA-117*; marine water - *Pelagibacter*, *Pelagilacustribacter*, *Actinomarina*, *CACPNW01*, *Marisimplicoccus*; phyllosphere - *Sphingomonas*, *Methylobacterium*, *Curtobacterium*, *Pantoea*, *Microbacterium*; soil - *Palsa-739*, *PALSA-600*, *JAQBQB01*, *JACVRW01*, *AC-32*; wastewater - *Aliarcobacter*, *Cloacibacterium*, *Acidovorax*, *Propionivibrio*, *Zoogloea* (Table S6).

The summed relative abundances of habitat indicator genera were always greater in their indicated habitats than in other habitats (Table 1, Figure 2). The median summed relative abundances of habitat indicator genera across a subset of 225 samples of the indicated habitat ranged from 11.2% to 89.4%, with the summed abundance being highest in human vaginal samples and lowest in freshwater sediment samples (Table 1). In general, spillover (i.e., indicator genera from one habitat found in other habitats) was minimal (median = 0, mean 0.07% to 1.26%), but there was some spillover among mammalian gut habitats and the three other human-associated habitats, as well as among soil and freshwater sediment and phyllosphere samples (Figure 2). Such spillover is unavoidable because most indicator genera are not exclusively found in the target habitat and there could also be errors in the metadata.

To demonstrate the utility of IndicPiper, we analyzed a depth profile and chronosequence of sediment samples that had accumulated over ∼2000 years in Lake Tjörnin, Iceland [15]. As expected, these samples were characterized mostly by indicators of freshwater sediment (51 genera) and water (39 genera) and marine sediment (19 genera) and water (22 genera) indicators, and to a lesser extent, glacier (7 genera), phyllosphere (3 genera), soil (61 genera), and wastewater (5 genera) indicators (Figure 4). Hydrothermal vent, human oral, and human skin indicator genera were also detected, but the mean summed relative abundances of these indicator genera were each < 0.15%. Lake Tjörnin used to be connected to the ocean > 1200 years ago and samples from that time period had appreciable marine water indicators (Figure 4). The lake could also be flooded at high tide during the 1700s to early 1900s, consistent with marine sediment taxa inputs, before being completely closed off by barriers in 1913 (Figure 4). Soil, phyllosphere, and glacier indicator taxa increased in relative abundance following human arrival around the year 800. Wastewater-associated taxa appeared in recent history following more intense and extensive urbanization around the lake (Figure 4).

**Figure 4.**
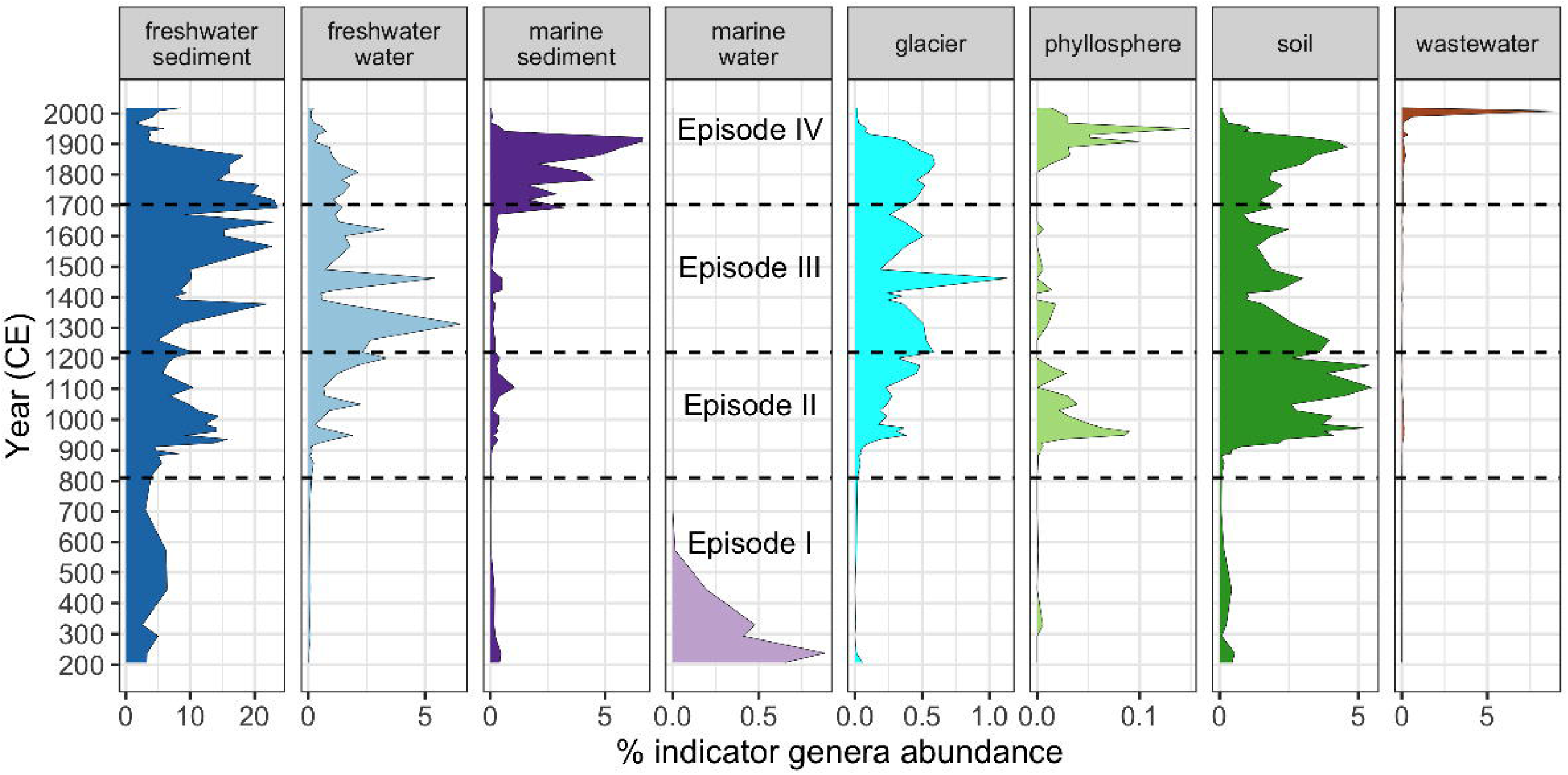
Summed relative abundances of habitat indicator genera over time in Lake Tjörnin, Iceland. The four episodes correspond to descriptions in Kjær et al. (2025). Note the difference in x-axis scale among panels. IndicPiper v1.0.0 was used for this analysis to match the GTDB r226 taxonomy used by Kjær et al. (2025). Note that the percent relative abundance is out of the total number bacterial and archaeal reads, not the entire metagenome.

As another demonstration of how IndicPiper could be used, we analyzed indicator genera in dust samples collected from the inside and outside of homes across the U.S [17]. These samples were characterized primarily by phyllosphere (10 genera), soil (152 genera), human skin (5 genera), human oral (34 genera), and mammalian gut (52 genera) indicator genera (Figure 4), with the summed relative abundance of phyllosphere indicator genera being the most abundant of any habitat (median ∼7%). Indicator genera of other habitats were detected but were less than 0.5% summed relative abundance on average. The summed relative abundances of phyllosphere indicator genera and soil indicator genera were each significantly greater in outdoor dust samples than in indoor dust samples (paired t-test p < 0.001, Figure 4). In contrast, the summed relative abundances of human skin, human oral, and mammalian gut indicator genera were each significantly greater in indoor dust samples than in outdoor dust samples (paired t-test, p < 0.001, Figure 5).

**Figure 5.**
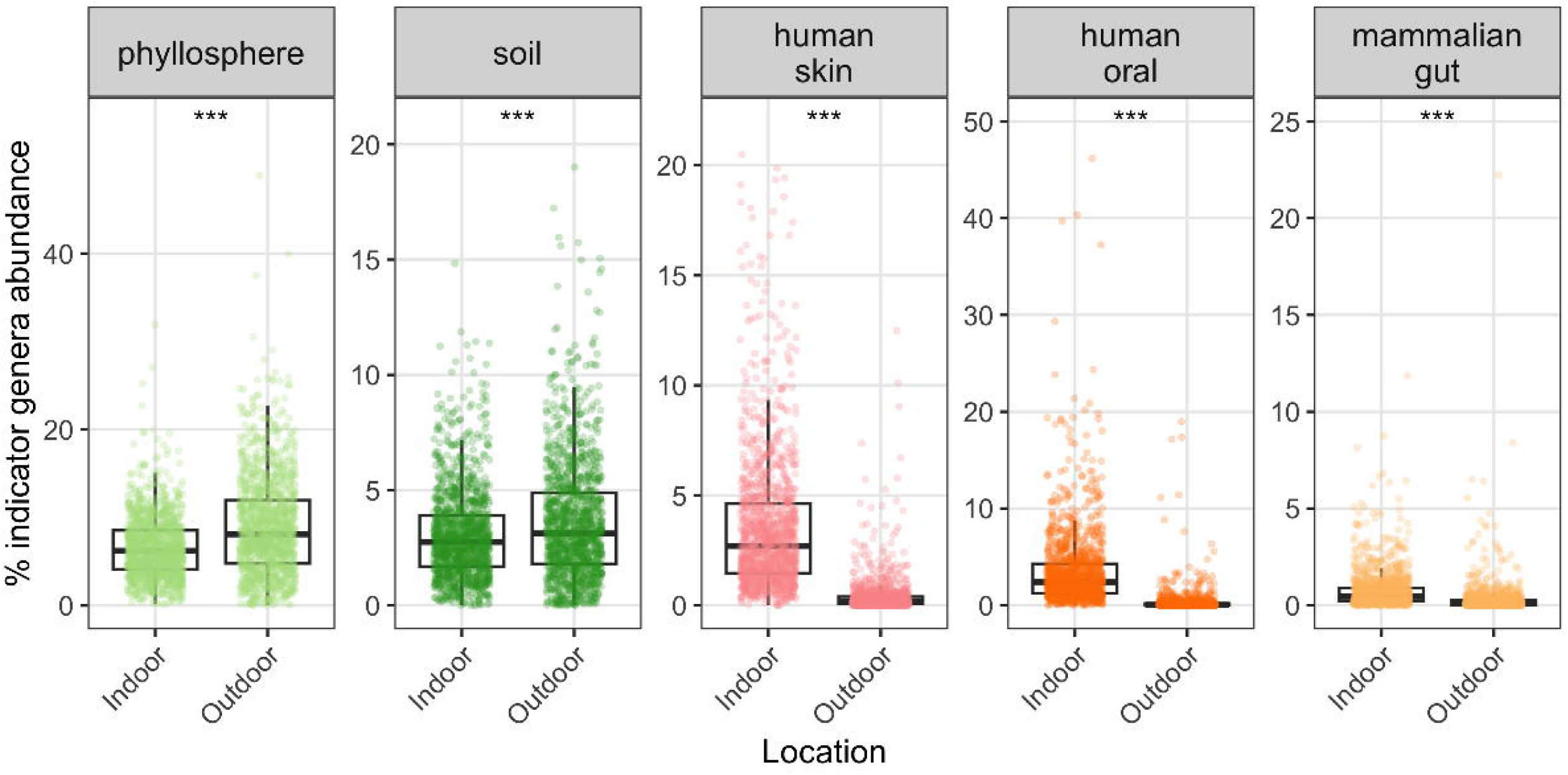
Summed relative abundances of habitat indicator genera in indoor and outdoor dust samples collected across the U.S (n = 1276 households with paired samples). *** = paired t-test p < 0.001. Note differences in the y-axis scale.

As a final example, we analyzed water samples from the Willamette River watershed in Oregon, USA [21] and found that they were characterized primarily by freshwater water (n = 30 genera), soil (n = 39 genera), glacier (n = 12 genera), freshwater sediment (n = 12 genera), and wastewater (n = 9 genera) indicator genera, with differences in relative abundance across the watershed from the mountains to the plains to urban areas (Figure 6). Hydrothermal vent, mammalian gut, marine water, marine sediment, and phyllosphere indicators were also detected, but the mean summed relative abundances of these indicator genera were all < 0.02%. Human oral, human skin, and human vaginal indicator genera were not detected. Freshwater water indicator genera had the greatest summed relative abundance in the plains, while freshwater sediment and soil indicator genera had the greatest summed relative abundance in the mountains (Figure 6). The summed relative abundances of glacier and wastewater indicator genera did not differ significantly across the three watershed zones (Figure 6).

**Figure 6.**
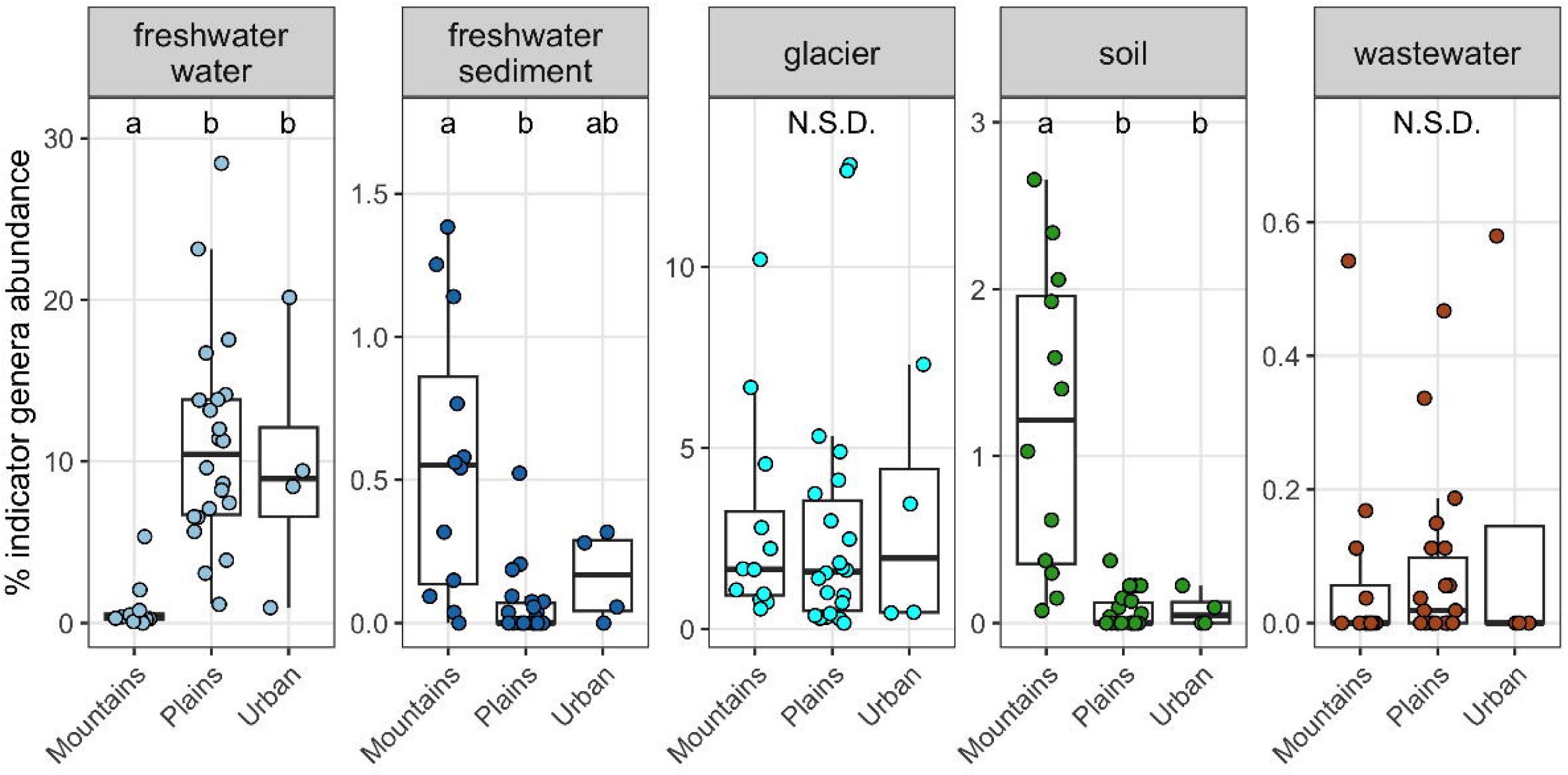
Summed relative abundances of habitat indicator genera in water samples collected from across the Willamette watershed in Oregon, USA, including mountains, plains, and urban areas. Different letters represent significant differences (Tukey’s posthoc, p < 0.05). N.S.D. = not significantly different. Note differences in the y-axis scale.

## Discussion

Here we present the IndicPiper methodology and database and demonstrate their utility for analyzing taxonomic profiles from microbial DNA sequencing data, including shotgun metagenomes and 16S rRNA amplicons. IndicPiper uses the GTDB taxonomic framework [11] and takes advantage of the large database of taxonomic profiles generated by SingleM that are housed in the Sandpiper database [6]. We note that users should assign taxonomy with GTDB, pay attention to which GTDB release they use, and use the corresponding IndicPiper versions (at the time of writing v1.0.0 = GTDB r226, v2.1.0 = GTDB r232), as genera can be added, deleted, or updated in each GTDB release. We plan on updating IndicPiper with each major Sandpiper/GTDB release, but this could also be done by any user by downloading the Sandpiper files from Zenodo, running the preliminary database filtering script (filterSandpiper.R), and then running the standard IndicPiper functions (Figure 1). For each release, users can use the provided database that was built using 13 habitats, which is likely applicable to many projects in microbial ecology. If users need to add or remove habitats to suit their research needs, they are encouraged to generate their own custom databases by supplying a list of habitats to the prepIndicPiper function and then executing the runIndicPiper function (Figure 1). For example, if there is only a subset of habitats of interest for a given study, identifying indicator taxa for just that subset of habitats is possible and may allow for more indicator genera to be identified, with the potential risk that those indicator genera may also be common in other habitats not considered.

We demonstrated IndicPiper on both shotgun metagenomic data (Lake Tjörnin) and 16S rRNA gene amplicon data (Willamette River watershed, household dust), and on three different sample types (sediment, dust, water). In all three examples, we could match a substantial proportion of the genera found in each dataset to the IndicPiper database, and IndicPiper made it possible to determine how taxa associated with different habitats changed over time or space in the given samples. Generally, the results were in line with expectations. Lake Tjörnin sediments were dominated by freshwater-associated taxa, the presence of marine-associated taxa corresponded to periods when the lake was connected to the ocean, and there was also evidence of recent human impact via the introduction of wastewater-associated taxa. In the household dust study, dust outside of homes had more phyllosphere- and soil-associated taxa, which are commonly found in airborne DNA samples [22–24], while dust inside of homes had more human-associated indicators. In the Willamette River watershed, soil and freshwater sediment indicators were relatively more abundant in the mountainous source waters and smaller order streams [25] than in the downstream waters. There were also some interesting and unexpected insights gained, such as a lack of an increase in wastewater taxa in urban areas of the Willamette River watershed (Figure 6). We use these datasets to demonstrate the utility of IndicPiper, but ultimately it is up to individual researchers to use IndicPiper for their own analyses and to interpret the IndicPiper results.

It is important to recognize that only a small subset of genera are considered indicators of certain habitats according to IndicPiper (880 genera in v1.0.0 and 767 genera in v2.1.0 out of more than 25,000 genera in the Sandpiper taxonomic profiles). Therefore, relative abundances of indicator taxa do not sum to 100% for any given community. Non-indicators are genera that could be prevalent in more than one habitat or genera that are not detected consistently enough in any one habitat type. For example, although *Arthrobacter* are common in many soil samples [26], this same genus is also prevalent in freshwater sediments so it is not considered an indicator of soil because the IndVal is < 0.5. These non-indicators will generally comprise a substantial proportion of the community of a given sample, and some habitats are likely to have a greater proportion of the community assigned as “non-indicator” (Figure 2). For example, phyllosphere, soil, freshwater sediment, freshwater water, marine sediment, hydrothermal vent, and wastewater habitats had median summed relative abundances of indicator genera of < 35% (Table 1), likely because they contain many taxa that are found in other environments or because the environments are very diverse and can have high compositional variation globally. In contrast, marine water, mammalian gut, and human-associated habitats (oral, skin, vaginal) have a much greater proportion of the community accounted for by indicator genera, likely because they are less variable and the taxa are more exclusive to those habitats. In the three case studies, the average relative abundance of non-indicator genera was 86% for Lake Tjörnin sediments, 35% for the household dust samples, and 17% for the Willamette River Watershed. Additionally, many sequences, particularly from short-read sequencing, cannot be confidently assigned to the genus level, and therefore cannot be matched with the IndicPiper database. For example, the average relative abundance of taxa with genus-level “NA” was 72% for the Willamette River watershed and 49% for the dust samples. For these reasons, when interpreting the results of IndicPiper applied to a given dataset, we recommend focusing on the relative differences in the summed abundances of indicators of a given habitat across time (Figure 4) or space (Figure 5, Figure 6), rather than focusing on the actual values of the summed abundances, except for general validation (i.e., more freshwater indicators in freshwater samples). This is also a key conceptual difference between IndicPiper and SourceTracker - for IndicPiper, a summed relative abundance of indicator genera for a particular habitat does not mean that 25% of all taxa in that sample came from that habitat. It simply means that there are genera that are significantly associated with a particular habitat, and the summed relative abundance of those particular genera is 25%.

It is also important to recognize the number and types of habitats selected for IndicPiper analysis will influence the genera that are determined to be indicators of each habitat. For example, if mammalian guts were not combined as done here and both human gut and primate gut were included in the selected habitats, many taxa that were mammalian gut indicators (significantly more prevalent in mammalian guts than other habitats) would likely be excluded, because taxa that are shared among human guts and primate guts would not be identified as indicators of either habitat.

Other limitations of IndicPiper include the standard limitations of metagenomic sequencing, including relative (rather than absolute) abundances and detection limits. We used a minimum cutoff of 150 million bacterial and archaeal bases [14] to remove metagenomes with poor sequencing depth of the prokaryotic community that may not have adequately captured the diversity of the sample. SingleM can detect rare taxa, as it reports the coverage and relative abundances for taxa with as low as 0.35x coverage. Taxa that are consistently below this threshold would likely be poor indicator taxa candidates. Furthermore, we removed samples with low complexity (samples with a single order making up > 95% of reads), so the IndicPiper input files did not contain any extremely high relative abundance values. Another benefit of SingleM with regards to the accuracy of the relative abundance profiles is its ability to identify and quantify novel lineages that currently lack genomic representation; SingleM outperformed other tools in this regard [6].

IndicPiper will continue to be improved as more archaeal and bacterial genomes are added to reference databases and classified to the genus level (with new GTDB releases) and as more metagenomes are profiled with SingleM (with new Sandpiper releases). IndicPiper can be updated following those releases to provide the most updated assessments of archaeal and bacterial habitat associations. The provided database for 13 habitats will likely be of use to many researchers, while those looking for more customization can use the provided functions (Figure 1) to generate their own databases with their own habitats of interest. The database could also be a starting point for other research questions in microbial ecology, such as investigating genomic attributes associated with indicator genera of different habitats, building on previous work on this topic [27–29]. Ultimately, IndicPiper takes advantage of pre-existing data and provides a tractable and straightforward approach to identify habitat associations of bacterial and archaeal genera, information that contributes to the ecological interpretation of microbial community sequencing data. We anticipate that the conceptual simplicity of IndicPiper and the utility of the results will be broadly helpful to microbial ecologists seeking to explore and glean new insights from their DNA sequencing datasets.

## Acknowledgements

This work was supported by The Ancient Environmental Genomics Initiative for Sustainability (AEGIS) project, which is supported by funding from the Novo Nordisk Foundation (NNF24SA0092560) and the Wellcome Trust (313162/Z/24/Z). The data generation infrastructure at the University of Copenhagen (used to generate the Lake Tjörnin data) is supported by funding from DNRF (DNRF174). BJW was supported by an Australian Research Council Future Fellowship (#FT210100521) and Discovery Project (#DP230101171). This work utilized the Alpine High-Performance Computing resource at the University of Colorado Boulder. Alpine is jointly funded by the University of Colorado Boulder, the University of Colorado Anschutz, Colorado State University, and the National Science Foundation (award 2201538).

## Data Availability

The Sandpiper database is available at https://zenodo.org/records/20437114. The starting metadata and taxonomic profile files for IndicPiper are available at https://zenodo.org/records/23424117. The IndicPiper code is available at https://github.com/cliffbueno/IndicPiper and the latest stable release is available at Zenodo (https://zenodo.org/records/23481965). The 16S rRNA gene taxonomy database for GTDB r232 is available at https://figshare.scilifelab.se/articles/dataset/SBDI_Sativa_curated_16S_GTDB_database/14869077.

## Notes

### Competing Interest Statement

The authors have declared no competing interest.

