## Supplementary material for "Comprehensive identification of bacterial and archaeal habitat associations using IndicPiper": Figure S2

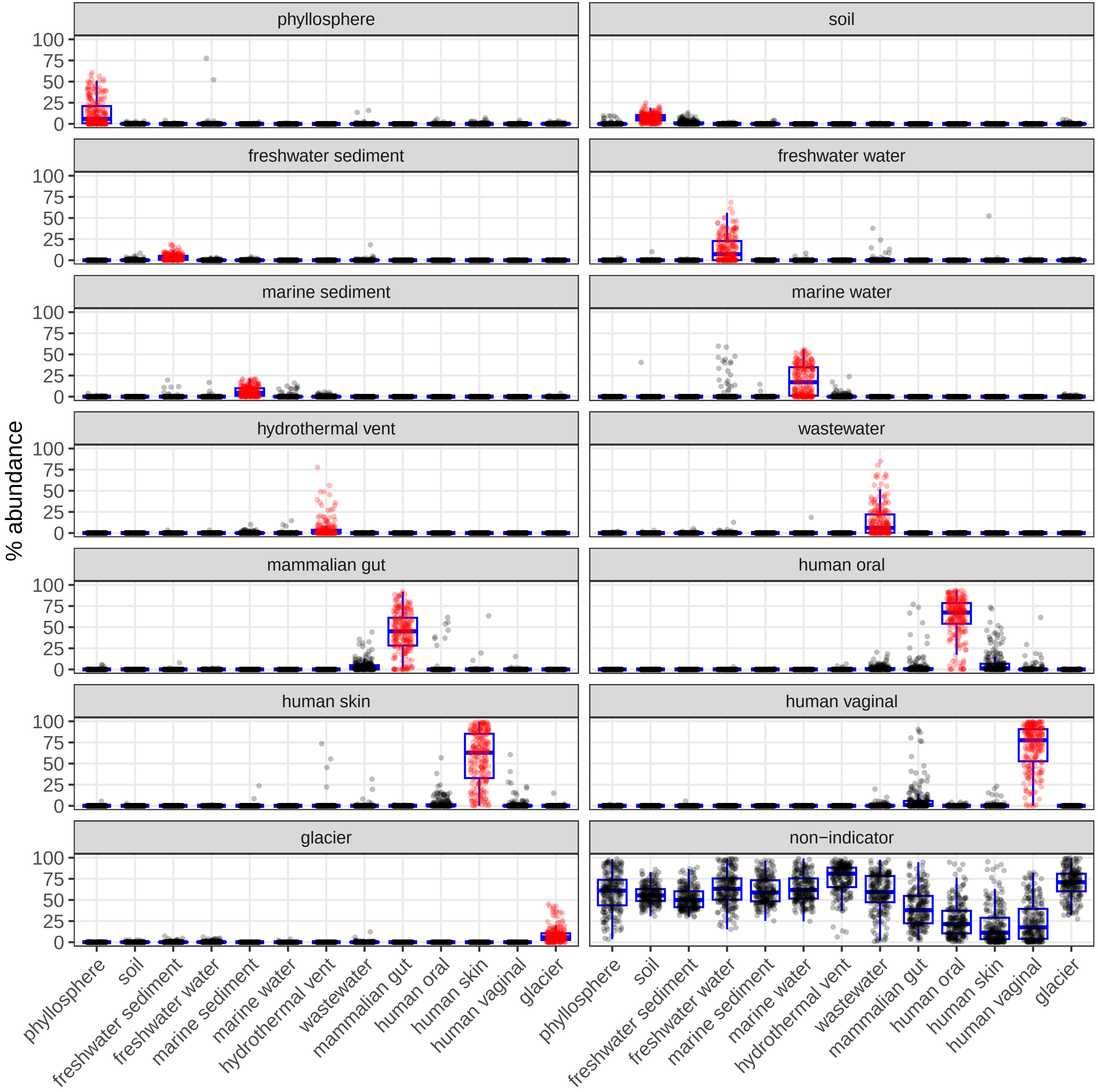

% abundance
phyllosphere
soil
freshwater sediment
freshwater water
marine sediment
marine water
hydrothermal vent
wastewater
mammalian gut
human oral
human skin
human vaginal
glacier
non−indicator
0
25
50
75
100
